# An *Escherichia coli* nitrogen starvation response is important for mutualistic coexistence with *Rhodopseudomonas palustris*

**DOI:** 10.1101/262139

**Authors:** Alexandra L. McCully, Megan G. Behringer, Jennifer R. Gliessman, Evgeny V. Pilipenko, Jeffrey L. Mazny, Michael Lynch, D. Allan Drummond, James B. McKinlay

## Abstract

Microbial mutualistic cross-feeding interactions are ubiquitous and can drive important community functions. Engaging in cross-feeding undoubtedly affects the physiology and metabolism of individual species involved. However, the nature in which an individual’s physiology is influenced by cross-feeding and the importance of those physiological changes for the mutualism have received little attention. We previously developed a genetically tractable coculture to study bacterial mutualisms. The coculture consists of fermentative *Escherichia coli* and phototrophic *Rhodopseudomonas palustris*. In this coculture, E. coli anaerobically ferments sugars into excreted organic acids as a carbon source for R. palustris. In return, a genetically-engineered R. palustris constitutively converts N_2_ into NH_4_^+^, providing *E. coli* with essential nitrogen. Using RNA-seq and proteomics, we identified transcript and protein levels that differ in each partner when grown in coculture versus monoculture. When in coculture with *R. palustris, E. coli* gene-expression changes resembled a nitrogen starvation response under the control of the transcriptional regulator NtrC. By genetically disrupting *E. coli* NtrC, we determined that a nitrogen starvation response is important for a stable coexistence, especially at low *R. palustris* NH_4_^+^ excretion levels. Destabilization of the nitrogen starvation regulatory network resulted in variable growth trends and in some cases, extinction. Our results highlight that alternative physiological states can be important for survival within cooperative cross-feeding relationships.

**Importance:** Mutualistic cross-feeding between microbes within multispecies communities is widespread. Studying how mutualistic interactions influence the physiology of each species involved is important for understanding how mutualisms function and persist in both natural and applied settings. Using a bacterial mutualism consisting of *Rhodopseudomonas palustris* and *Escherichia coli* growing cooperatively through bidirectional nutrient exchange, we determined that an *E. coli* nitrogen starvation response is important for maintaining a stable coexistence. The lack of an *E. coli* nitrogen starvation response ultimately destabilized the mutualism and, in some cases, led to community collapse after serial transfers. Our findings thus inform on the potential necessity of an alternative physiological state for mutualistic coexistence with another species compared to the physiology of species grown in isolation.

## Introduction

Within diverse microbial communities, species engage in nutrient cross-feeding with reciprocating partners as a survival strategy (1). In cases where species are not obligate mutualists, transitioning from a free-living lifestyle to one based on cross-feeding can change the physiological state of the cells involved, the extent to which depends on the nature of the cross-feeding relationship. For example, cross-feeding can promote physiological changes that increase virulence (2, 3) or drastically alter cellular metabolism (4), in some cases allowing for lifestyles that are only possible during mutualistic growth with a partner (4–7). Aside from these examples, relatively little is known about how cell physiology is influenced by mutualistic cross-feeding, despite the prevalence of cross-feeding in microbial communities.

Synthetic communities, or cocultures, are ideally suited for studying the physiological responses to cooperative cross-feeding given their tractability (8, 9). We previously developed a bacterial coculture that consists of fermentative *Escherichia coli* and the N_2_-fixing photoheterotroph *Rhodopseudomonas palustris* (Fig. 1) (10). In this coculture, *E. coli* anaerobically ferments glucose into organic acids, providing *R. palustris* with essential carbon. In return, a genetically engineered *R. palustris* strain (Nx) constitutively fixes N_2_ gas, resulting in NH_4_^+^ excretion that provides *E. coli* with essential nitrogen. The result is an obligate mutualism that maintains a stable coexistence and reproducible growth trends (10) as long as bidirectional nutrient cross-feeding levels are maintained within a defined range (11, 12).

Here we determined how nutrient cross-feeding between *E. coli* and *R. palustris* Nx alters the physiological state of each partner population. Using RNA-seq and proteomic analyses, we identified genes in both species that were differentially expressed in coculture compared to monoculture, with *E. coli* exhibiting more overall changes in gene expression than *R. palustris* Nx. Specifically, *E. coli* gene-expression patterns resembled that of nitrogen-deprived cells, as many upregulated genes were within the nitrogen-starvation response regulon, controlled by the master transcriptional regulator NtrC. Genetic disruption of *E. coli ntrC* resulted in variable growth trends at low *R. palustris* NH_4_^+^ excretion levels and prevented long-term mutualistic coexistence with *R. palustris* across serial transfers. Our results highlight the fact that cross-feeding relationships can stimulate alternative physiological states for at least one of the partners involved and that adjusting cell physiology to these alternative states can be critical for maintaining coexistence.

## Results

### Engaging in an obligate mutualism alters the physiology of cooperating partners

In our coculture, *E. coli* and *R. palustris* Nx carry out complementary anaerobic metabolic processes whose products serve as essential nutrients for the respective partner. Specifically, *E. coli* ferments glucose into acetate, lactate, and succinate, which serve as carbon sources for *R. palustris* Nx, while other fermentation products such as formate and ethanol accumulate; in return *R. palustris* Nx fixes N_2_ and excretes NH_4_^+^ as the nitrogen source for *E. coli* (Fig. 1). We previously demonstrated that our coculture supports a stable coexistence and exhibits reproducible growth and metabolic trends when started from a wide range of starting species ratios, including single colonies (10). However, we hypothesized that coculture conditions would affect the physiology of each species, particularly *E. coli*, based on the following observations. First, as growth is coupled in our coculture, *E. coli* is forced to grow 4.6-times slower in coculture with *R. palustris* Nx than it does in monoculture with abundant NH_4_^+^ due to slow NH_4_^+^ cross-feeding from *R. palustris* Nx (10). In contrast, *R. palustris* Nx grows at a rate in coculture that is comparable to that in monoculture (12), consuming a mixed pool of excreted organic acids from *E. coli*. Second, coculturing pulls *E. coli* fermentation forward due to removal of inhibitory end products. For example, we observed higher yields of formate, an *E. coli* fermentation product that *R. palustris* does not consume, in cocultures compared to *E. coli* monocultures (10).

**FIG 1.**
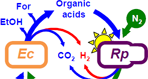
Bidirectional cross-feeding of carbon and nitrogen in an anaerobio bacterial mutualism between fermentative *Escherichia coli (Ec)* and phototrophic *Rhodopseudomonas palustris (Rp)*. *E. coli* anaerobically ferments glucose into excreted organic acids that *R. palustris* Nx consumes (acetate, lactate and succinate) and other products that *R. palustris* Nx does not consume (formate (For) and ethanol (EtOH)). In return, *R. palustris* Nx constitutively fixes N_2_ gas and excretes NH_4_^+^, supplying *E. coli* with essential nitrogen. *R. palustris* Nx grows photoheterotrophically wherein organic compounds are used for carbon and electrons, and light is used for energy.

To determine changes in gene-expression patterns imposed by coculturing, we performed RNA-seq and comparative proteomic analyses (13) on exponential phase cocultures and monocultures of *E. coli* and *R. palustris* Nx. To make direct comparisons, all cultures were grown in the same basal anaerobic minimal medium, and monocultures were supplemented with the required carbon or nitrogen sources to permit growth for each species. Cocultures and *E. coli* monocultures were provided glucose as a sole carbon source, whereas a mixture of organic acids and bicarbonate was provided to *R. palustris* Nx monocultures, as *R. palustris* does not consume glucose. For a nitrogen source, all cultures were grown under a N_2_ headspace, and *E. coli* monocultures were further supplemented with NH_4_Cl, as *E. coli* is incapable of using N_2_. We identified several differentially expressed genes between monoculture and coculture conditions in both species with more differences observed in *E. coli* compared to *R. palustris* Nx, in agreement with our initial hypothesis (Fig. 2). For *E. coli*, out of 4377 ORFs, 55 were upregulated and **68** were downregulated (Table 1) (log2 value cutoff=2). Out of 4836 ORFs in *R. palustris* Nx, 14 were upregulated and 20 were downregulated (Table 1) (log2 value cutoff=2). We also considered that due to lower *E. coli* abundance in coculture, the apparently larger *E. coli* gene response may be partly due to decreased resolution and thus increased error variance. Reassuringly, many of the genes identified as being differentially expressed by RNA-seq were in agreement with the proteomic results (Table 2). Both RNA-seq and proteomic analyses identified the *E. coli* ammonium transporter AmtB as an important, upregulated gene in coculture, corroborating our previous findings that *E. coli* AmtB activity is important for stable coexistence with *R. palustris* (12). Many *E. coli* genes involved in amino acid and purine biosynthesis were downregulated in coculture (Table 1, Table 2), consistent with the lower observed growth rate. Additionally, many *E. coli* flagellar and chemotaxis proteins were downregulated in coculture (Table 1, Table 2), perhaps suggesting that motility is not important for coculture growth. Alternatively, lower flagellar and chemotaxis transcript levels could be part of a general stress response (14), perhaps associated with nitrogen limitation in cocultures. Whereas many of the differentially expressed *E. coli* genes have been characterized in the literature, the *R. palustris* genes showing the largest differential expression were uncharacterized genes encoding upregulated putative alcohol/aldehyde dehydrogenases and a downregulated putative TonB-dependent receptor/siderophore (Table 1, Table 2). Together, these datasets provide insight on how engaging in obligate cross-feeding changes the lifestyle of each partner.

**FIG 2.**
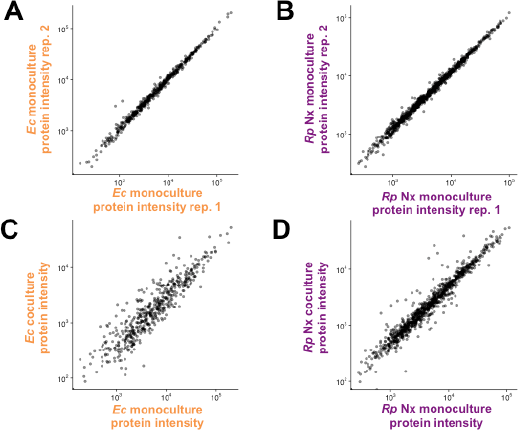
Coculture conditions result in altered protein expression patterns in both species, with more differences in WT *E. coli* compared to *R. palustris* Nx. Protein expression (estimated by LC-MS/MS intensity) of wild-type *E. coli* (left, **A,C**) and *R. palustris* Nx (right, **B, D**) comparing protein expression patterns between monoculture biological replicates (rep. 1 versus rep. 2, **A, B**) and monoculture (average over monoculture replicates) versus coculture (**C, D**).

**Table 1.**
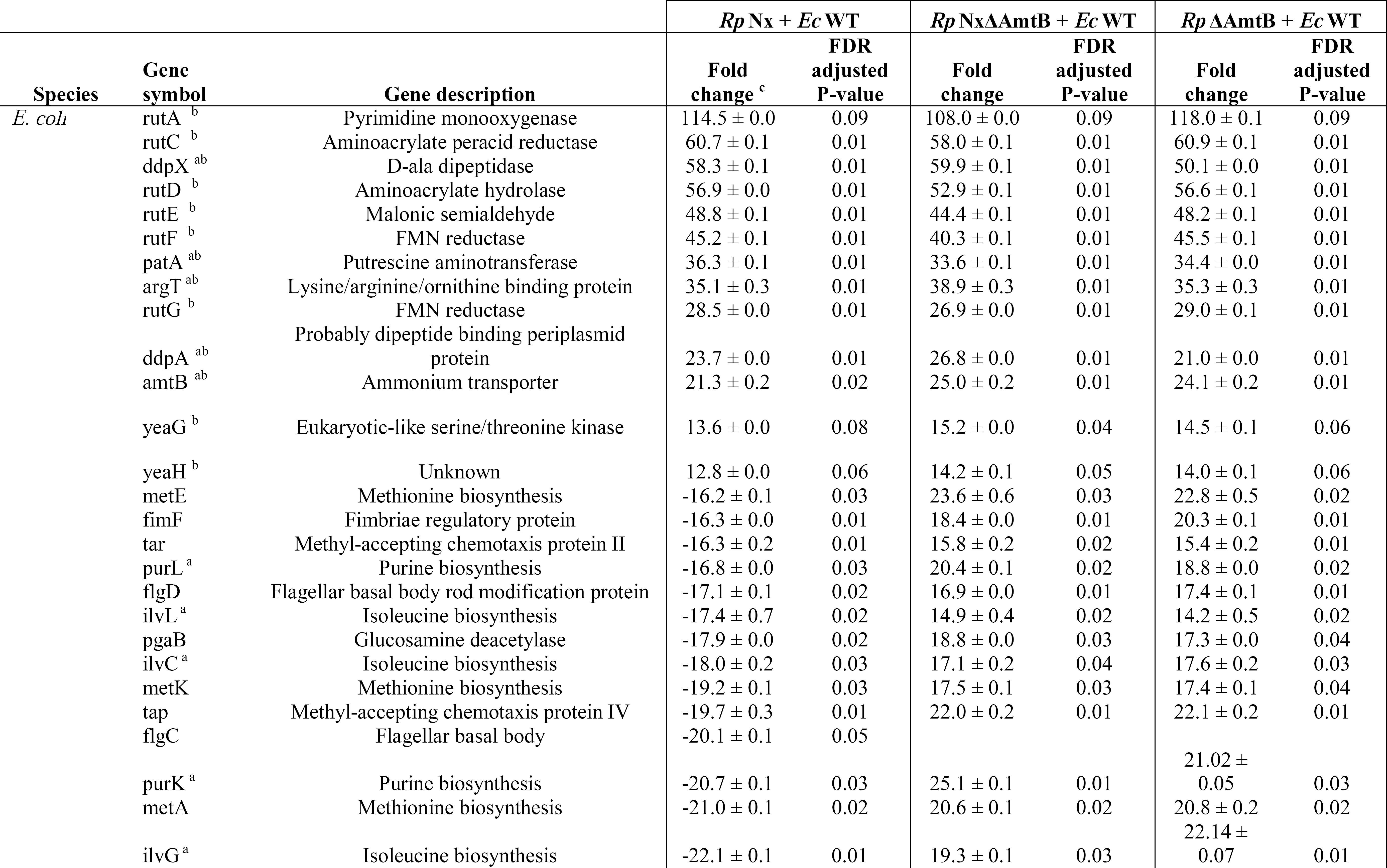
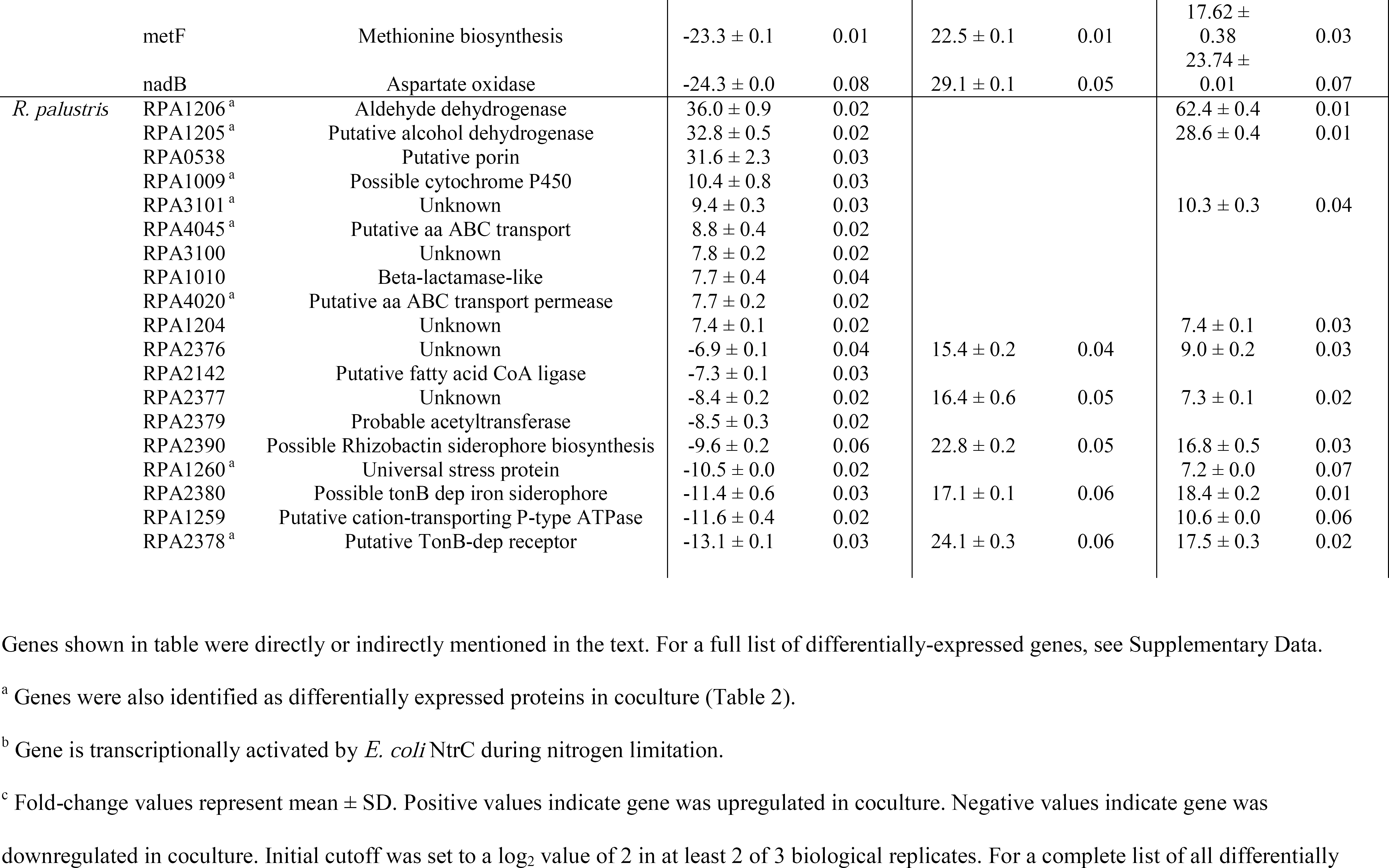
Selected differentially expressed transcripts in cocultures of *E. coli* and *R. palustris* compared to monocultures.

**Table 2.**
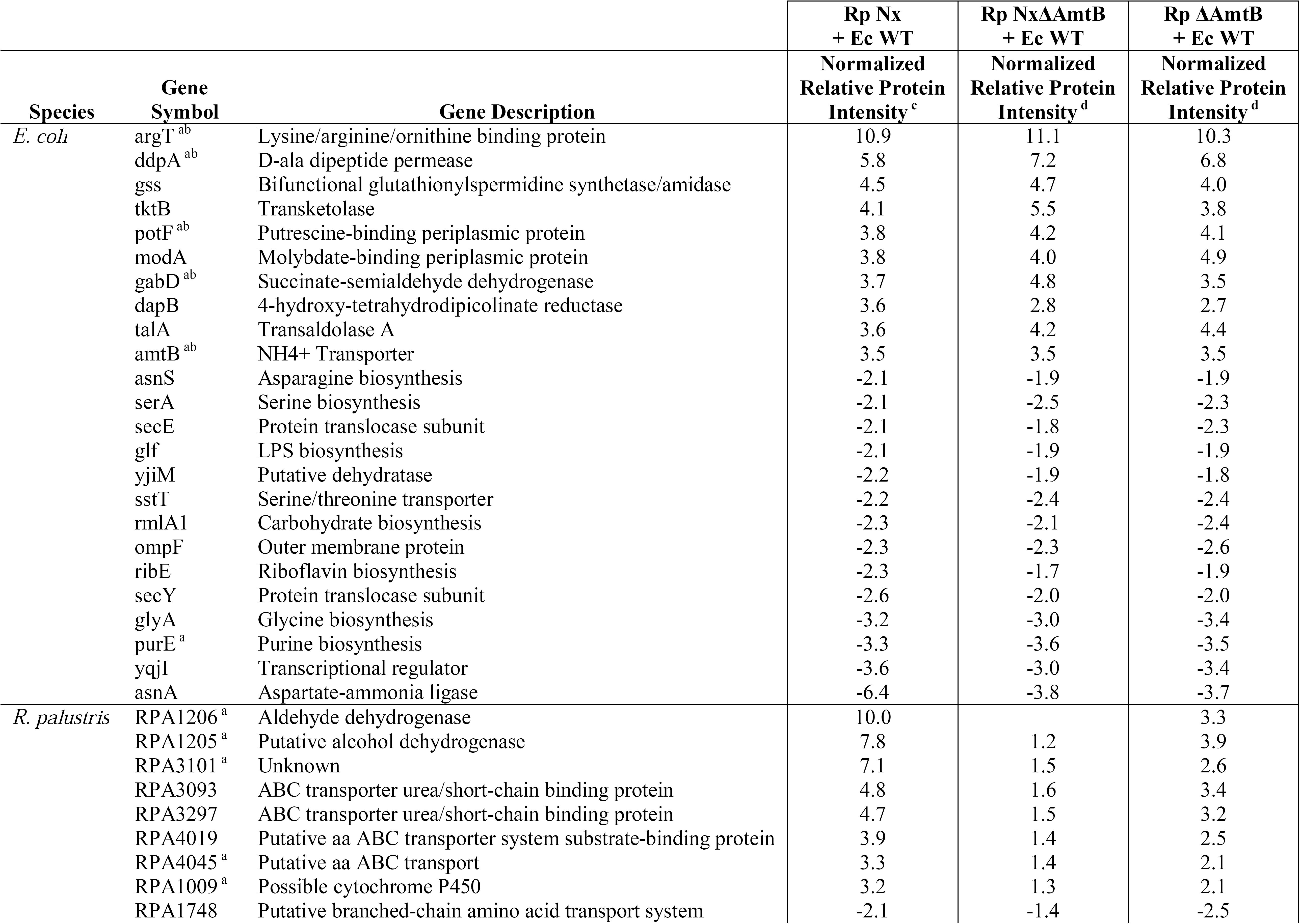
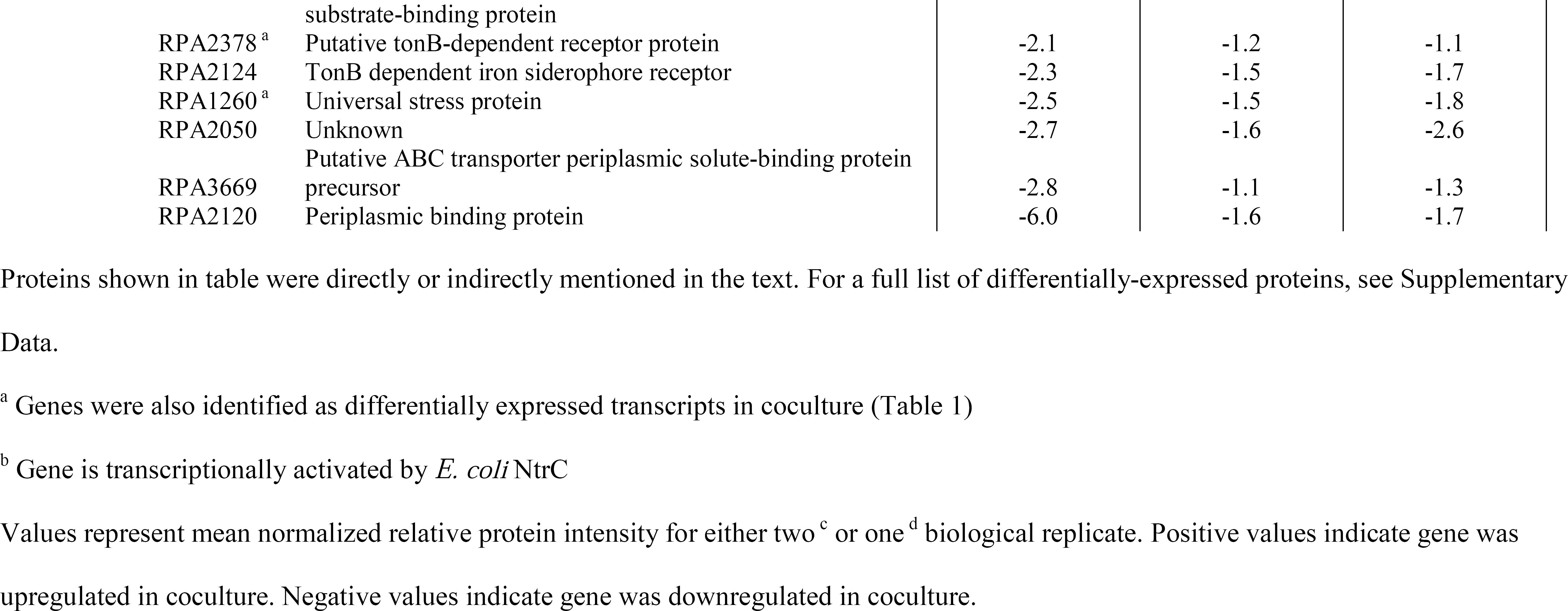
Selected differentially expressed proteins in cocultures of *E. coli* and *R. palustris* compared to monocultures.

| Species | Gene Symbol | Gene Description | Rp Nx<br>+ Ec WT | Rp NxΔAmtB<br>+ Ec WT | Rp ΔAmtB<br>+ Ec WT |
| --- | --- | --- | --- | --- | --- |
|  |  |  | Normalized<br>Relative Protein<br>Intensity<sup>c</sup> | Normalized<br>Relative Protein<br>Intensity<sup>d</sup> | Normalized<br>Relative Protein<br>Intensity<sup>d</sup> |
| *E. coli* | argT<sup>ab</sup> | Lysine/arginine/ornithine binding protein | 10.9 | 11.1 | 10.3 |
|  | ddpA<sup>ab</sup> | D-ala dipeptide permease | 5.8 | 7.2 | 6.8 |
|  | gss | Bifunctional glutathionylspermidine synthetase/amidase | 4.5 | 4.7 | 4.0 |
|  | tktB | Transketolase | 4.1 | 5.5 | 3.8 |
|  | potF<sup>ab</sup> | Putrescine-binding periplasmic protein | 3.8 | 4.2 | 4.1 |
|  | modA | Molybdate-binding periplasmic protein | 3.8 | 4.0 | 4.9 |
|  | gabD<sup>ab</sup> | Succinate-semialdehyde dehydrogenase | 3.7 | 4.8 | 3.5 |
|  | dapB | 4-hydroxy-tetrahydrodipicolinate reductase | 3.6 | 2.8 | 2.7 |
|  | talA | Transaldolase A | 3.6 | 4.2 | 4.4 |
|  | amtB<sup>ab</sup> | NH<sub>4</sub><sup>+</sup> Transporter | 3.5 | 3.5 | 3.5 |
|  | asnS | Asparagine biosynthesis | -2.1 | -1.9 | -1.9 |
|  | serA | Serine biosynthesis | -2.1 | -2.5 | -2.3 |
|  | secE | Protein translocase subunit | -2.1 | -1.8 | -2.3 |
|  | glf | LPS biosynthesis | -2.1 | -1.9 | -1.9 |
|  | yjiM | Putative dehydratase | -2.2 | -1.9 | -1.8 |
|  | sstT | Serine/threonine transporter | -2.2 | -2.4 | -2.4 |
|  | rmlA1 | Carbohydrate biosynthesis | -2.3 | -2.1 | -2.4 |
|  | ompF | Outer membrane protein | -2.3 | -2.3 | -2.6 |
|  | ribE | Riboflavin biosynthesis | -2.3 | -1.7 | -1.9 |
|  | secY | Protein translocase subunit | -2.6 | -2.0 | -2.0 |
|  | glyA | Glycine biosynthesis | -3.2 | -3.0 | -3.4 |
|  | purE<sup>a</sup> | Purine biosynthesis | -3.3 | -3.6 | -3.5 |
|  | yqjI | Transcriptional regulator | -3.6 | -3.0 | -3.4 |
|  | asnA | Aspartate-ammonia ligase | -6.4 | -3.8 | -3.7 |
| *R. palustris* | RPA1206<sup>a</sup> | Aldehyde dehydrogenase | 10.0 |  | 3.3 |
|  | RPA1205<sup>a</sup> | Putative alcohol dehydrogenase | 7.8 | 1.2 | 3.9 |
|  | RPA3101<sup>a</sup> | Unknown | 7.1 | 1.5 | 2.6 |
|  | RPA3093 | ABC transporter urea/short-chain binding protein | 4.8 | 1.6 | 3.4 |
|  | RPA3297 | ABC transporter urea/short-chain binding protein | 4.7 | 1.5 | 3.2 |
|  | RPA4019 | Putative aa ABC transporter system substrate-binding protein | 3.9 | 1.4 | 2.5 |
|  | RPA4045<sup>a</sup> | Putative aa ABC transport | 3.3 | 1.4 | 2.1 |
|  | RPA1009<sup>a</sup> | Possible cytochrome P450 | 3.2 | 1.3 | 2.1 |
|  | RPA1748 | Putative branched-chain amino acid transport system | -2.1 | -1.4 | -2.5 |

|  |  |  |  |  |
| --- | --- | --- | --- | --- |
|  | substrate-binding protein |  |  |  |
| RPA2378<sup>a</sup> | Putative tonB-dependent receptor protein | -2.1 | -1.2 | -1.1 |
| RPA2124 | TonB dependent iron siderophore receptor | -2.3 | -1.5 | -1.7 |
| RPA1260<sup>a</sup> | Universal stress protein | -2.5 | -1.5 | -1.8 |
| RPA2050 | Unknown | -2.7 | -1.6 | -2.6 |
|  | Putative ABC transporter periplasmic solute-binding protein |  |  |  |
| RPA3669 | precursor | -2.8 | -1.1 | -1.3 |
| RPA2120 | Periplasmic binding protein | -6.0 | -1.6 | -1.7 |

Proteins shown in table were directly or indirectly mentioned in the text. For a full list of differentially-expressed proteins, see Supplementary Data.
<sup>a</sup> Genes were also identified as differentially expressed transcripts in coculture (Table 1)
<sup>b</sup> Gene is transcriptionally activated by *E. coli* NtrC
Values represent mean normalized relative protein intensity for either two<sup>c</sup> or one<sup>d</sup> biological replicate. Positive values indicate gene was upregulated in coculture. Negative values indicate gene was downregulated in coculture.

### An *E. coli* nitrogen starvation response is important for mutualistic growth with *R. palustris*

We chose to further examine differential gene expression patterns in *E. coli* as its growth rate and fermentation profile are drastically affected by coculturing, whereas the *R. palustris* Nx growth rate is similar to that in monoculture. We identified several *E. coli* genes and proteins that were upregulated in coculture with *R. palustris* Nx compared to monoculture growth (Table 1, Table 2). We hypothesized that the deletion of highly upregulated *E. coli* genes would negatively affect its growth in coculture. We made deletions in *E. coli* genes that were identified in both RNA-seq and proteome datasets as well as the highest upregulated *E. coli* transcript *(rutA)*. We did not examine the effect of deleting *amtB* in this case as we previously determined it to be important for coculture growth (12). These selected *E. coli* genes were all involved in metabolism of alternative nitrogen sources such as D-ala-D-ala dipeptides *(ddpX, ddpA)* (15), pyrimidines (rutA) (16), amino acids *(argT)* (17), and polyamines *(patA*, *potF)* (18). In monocultures with 15mM NH_4_C1, there were negligible differences in growth or fermentation profiles between WT *E. coli* and any of the single deletion mutants (Fig. S1). These results are consistent with findings by others, as these genes are only important when scavenging alternative nitrogen sources that are not present in our defined medium. We next tested these *E. coli* mutants in coculture with *R. palustris* Nx to determine if these genes were important when NH_4_^+^ is slowly cross-fed from *R. palustris* Nx. All cocultures using the *E. coli* mutants paired with *R. palustris* Nx exhibited similar growth and population trends to cocultures with WT *E. coli* (Fig. 3). Additionally, there were no significant differences in the growth rates, growth yields, or product yields from cocultures containing the *E. coli* mutants (Fig. S2). These data suggest that none of these highly expressed *E. coli* genes are solely important for coculture growth. While it is possible that synergistic expression of these genes is important for *E. coli’s* lifestyle in coculture, the actual nitrogen sources accessed by expression of these genes are absent in the defined medium. Thus, unless *E. coli* gains access to alternative nitrogen sources that we are unaware of in coculture with *R. palustris* Nx, synergistic expression of these genes likely provides little to no benefit.

**FIG S1.**
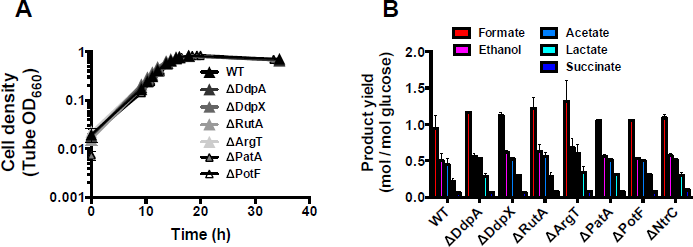
Single deletions of *E. coli* genes that were upregulated in coculture no effect in monoculture with 15 mM NH_4_^+^. Growth curves (**A**) and product yields (**B**) from *E. coli* monocultures grown with 15 mM NH_4_Cl. Product yields were taken in stationary phase. Error bars indicate SD, n=3.

**FIG S2.**
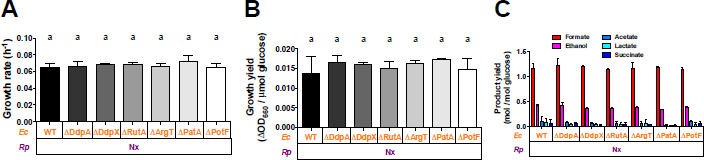
Additional trends from cocultures pairing *R. palustris* Nx with *E. coli* single deletions mutants. Growth rates (**A**), growth yields (**B**), and product yields (**C**) after a one-week culturing period from cocultures pairing *E. coli* mutants with deletions in highly upregulated genes with *R. palustris* Nx. Growth and product yields were taken at the final time point indicated in Fig. 3A. Cocultures were started with a 1% inoculum of stationary starter cocultures grown from single colonies. Error bars indicate SD, n=3. Different letters indicate statistical differences, p < 0.05, determined by one-way ANOVA with Tukey’s multiple comparisons posttest.

**FIG 3.**
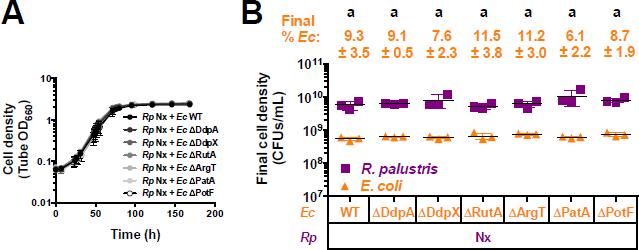
Single deletions of upregulated *E. coli* genes do not impair mutualistic growth with *R*. palustris Nx. Growth curves (**A**) and final cell densities (**B**) from cocultures pairing *E. coli (Ec)* mutants with deletions in highly upregulated genes with *R. palustris (Rp)* Nx. Final cell densities (**B**) were taken at the final time point in (**A**). Cocultures were started with a 1% inoculum of stationary starter cocultures grown from single colonies. Error bars indicate SD, n=3. Different letters indicate statistical differences, p < 0.05, determined by one-way ANOVA with Tukey’s multiple comparisons posttest.

Even though individual deletions of the *E. coli* genes showing high expression in coculture had no effect on coculture trends, we noted that they were all involved in nitrogen scavenging and fell within the regulon of the transcription factor, NtrC, which controls the nitrogen starvation response (19). During nitrogen limitation, the sensor kinase NtrB phosphorylates the response regulator NtrC (19). Phosphorylated NtrC then binds to DNA and activates expression of ∼45 genes (20), including those we tested genetically above and *amtB*, which we previously determined to be important for coculture growth (12). To examine the importance of the *E. coli* nitrogen starvation response in coculture, we deleted *ntrC*. We first checked for any general defects of the resulting ΔNtrC mutant in monoculture with 15 mM NH_4_Cl and found that it exhibited similar growth and metabolic trends to WT *E. coli* (Fig. S3). We then paired *E. coli* ΔNtrC with *R. palustris* Nx in coculture. Compared to cocultures using WT *E. coli*, cocultures with *E. coli* ΔNtrC exhibited slower growth rates, longer lag periods (Fig. 4A), and lower final *E. coli* cell densities (Fig. 4D). The long lag phase was less prominent in cocultures inoculated from single colonies (Fig. S4A) compared to cocultures inoculated with a 1% dilution of stationary cocultures (Fig. 4A). This result suggests that starting *E. coli* ΔNtrC cocultures from single colonies stimulated early growth, perhaps by increasing the *E. coli* frequency to be similar to that of *R. palustris* when started with colonies of similar sizes rather than a dilution of stationary cocultures wherein the *E. coli* frequency was low (∼0.1%; Fig. 4D). A higher initial *E. coli* frequency might help *E. coli* acquire excreted NH_4_^+^ before it is taken back up by *R. palustris* cells and thereby promote reciprocal cross-feeding, similar to what we observed previously in cocultures with *E. coli* ΔAmtB mutants that were defective for NH_4_^+^ uptake (12).

**FIG S3.**
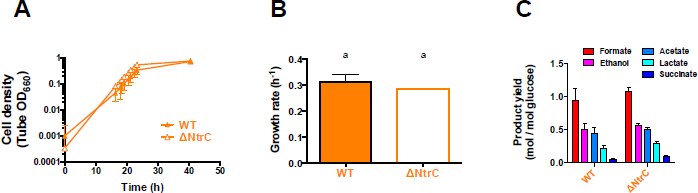
*E. coli* ΔNtrC growth and metabolic trends are similar to those of WT *E. coli* in monoculture with 15 mM NH_4_^+^. Growth curves (**A**), growth rate (**B**) and product yields (**C**) from WT *E. coli* (filled) or ΔNtrC (open) monocultures grown with 15 mM NH_4_CL Product yields were taken in stationary phase. Error bars indicate SD, n=3.

**FIG S4.**
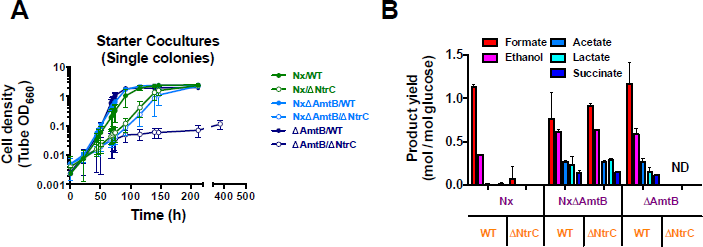
Additional trends from cocultures of E. coli ΔNtrC paired with different R. palustris Partners. Growth curves of starter cocultures inoculated with single colonies of each species (**A**) and product yields from test cocultures (**B**). Product yields (**B**) were taken at the final time point indicated in the respective growth curve in Fig. 4. Test cocultures were started with a 1% inoculum of stationary starter cocultures. Error bars indicate SD, n=3. ND, not determined.

**FIG 4.**
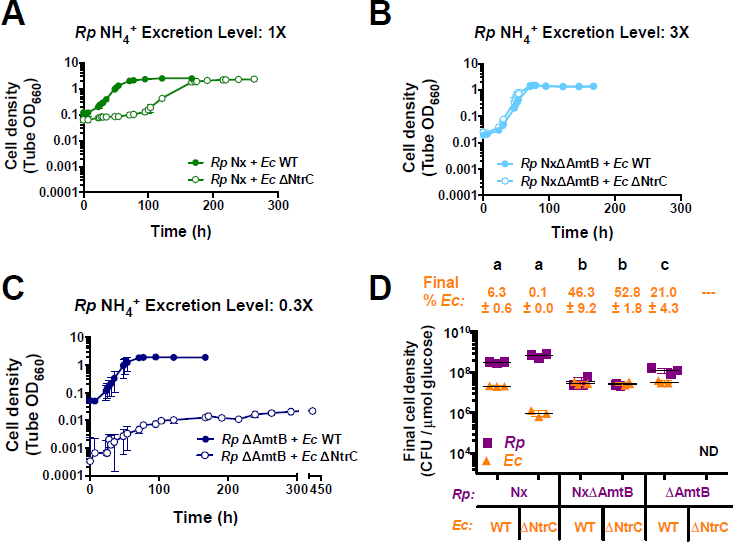
R. palustris NH_4_+ excretion level affects growth and population trends in cocultures with *E. coli* NtrC. Growth curves (**A, B, C**) and final cell densities normalized to glucose consumption (**D**) from cocultures pairing WT *E. coli (Ec)* (filled circles) or ΔNtrC (open circles) with *R. palustris (Rp)* strains with different NH_4_^+^ excretion levels. Final cell densities (**D**) were taken at the final time point in the respective growth curve (**A, B, C**), except for cocultures pairing *R. palustris* ΔAmtB with *E. coli* ΔNtrC which were sampled at 260 h. Cell densities were normalized to glucose consumed to account for incomplete glucose consumption in cocultures containing *E. coli* ΔNtrC. Cocultures were started with a 1% inoculum of stationary starter cocultures grown from single colonies. Error bars indicate SD, n=3. Different letters indicate statistical differences, p < 0.05, determined by one-way ANOVA with Tukey’s multiple comparisons posttest. ND, not determined.

The overall coculture metabolism was also altered when *E. coli* ΔNtrC was paired with *R. palustris* Nx. In cocultures pairing WT *E. coli* with *R. palustris* Nx, glucose is typically fully consumed within 5 days coinciding with the accumulation of formate and ethanol (10). Cocultures pairing *E. coli* ΔNtrC with *R. palustris* Nx differed in this regard, leaving ∼40% of the glucose unconsumed after 10 days and exhibiting little to no formate and ethanol accumulation (Fig. S4B). Even despite the lower glucose consumption, the final *R. palustris* cell density of cocultures pairing *R. palustris* Nx with *E. coli* ΔNtrC was similar to those with WT *E. coli*. This unexpectedly high cell density could be explained by consumption of formate and ethanol by *R. palustris* Nx, though we have never observed consumption of formate by *R. palustris* Nx in monoculture. Alternatively, a lack of formate and/or ethanol production by *E. coli* could explain the high cell density if the fermentation profile were shifted towards organic acids that *R. palustris* normally consumes, namely acetate, lactate and succinate. Together, these data indicate that misregulation of the nitrogen starvation response affected coculture growth and metabolism.

As noted above, the low *E. coli* ΔNtrC population and decreased coculture growth rate when paired with *R. palustris* Nx resembled trends from cocultures that contained *E. coli* ΔAmtB mutants (12). We previously found that the *E. coli* NH_4_^+^ transporter, AmtB, was required for coexistence with *R. palustris* Nx across serial transfers as the transporter gives *E. coli* a competitive advantage in acquiring the transiently available NH_4_^+^ before it can be reclaimed by the *R. palustris* population (12). To determine if *E. coli* ΔNtrC was capable of maintaining a stable coexistence in coculture, we inoculated cocultures of *E. coli* ΔNtrC paired with *R. palustris* Nx at equivalent CFUs and performed serial transfers every 10 days. While average final *E. coli* frequencies were consistently between 0.6 – 2.8 % (Fig. 5A), the values became variable over serial transfers, as did coculture growth rates, lag periods, and net changes in both *E. coli* and *R. palustris* cell densities (Fig. 5). This variability was due to 2 of the 4 lineages exhibiting improved coculture growth over successive transfers Fig. 5B, C), perhaps due to the emergence of compensatory mutations, while the other two lineages showed declining growth trends (Fig. 5D, E). Indeed, by transfers 5 and 6 there was little to no coculture growth in the slower-growing lineages (Fig 4D, E). The heterogeneity in growth trends through serial transfers of cocultures with *E. coli* ΔNtrC is in stark contrast to the stability of cocultures with WT *E. coli*, which we have serially transferred over 100 times with no extinction events (McKinlay, unpublished data). The nitrogen starvation response thus appears to be important for long-term survival of the mutualism.

**FIG 5.**
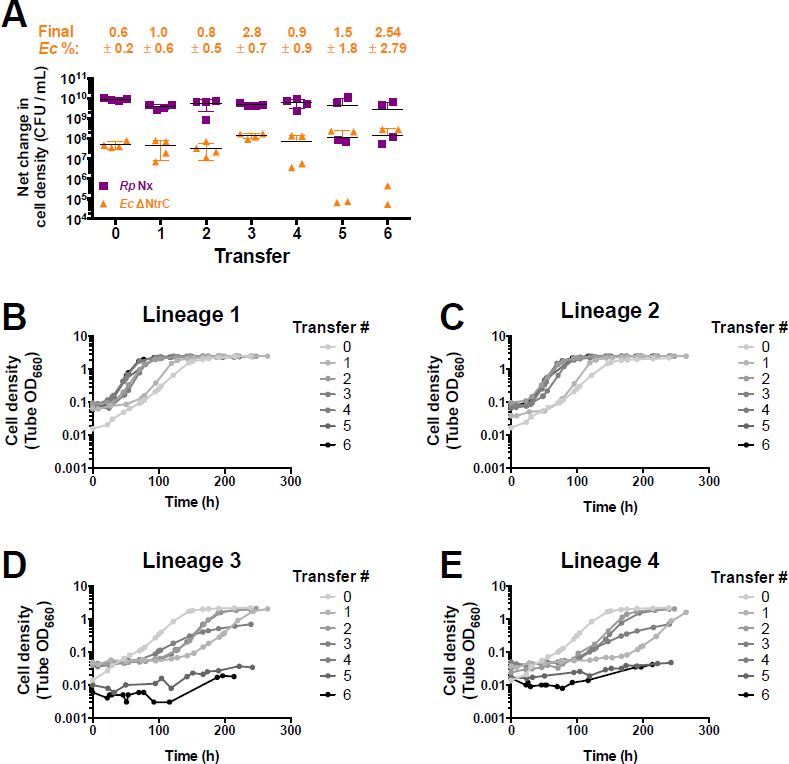
Lack of *E. coli* NtrC results in variable coculture growth trends across serial transfers. Net changes in cell densities (**A**) and replícate growth curves (**B-E**) of cocultures pairing *E. coli (Ec)* ΔNtrC with *R. palustris (Rp)* Nx across serial transfers. Cocultures were initially inoculated (Transfer 0) at a 1:1 starting species ratios based on CFUs/mL from *R. palustris* and *E. coli* monocultures. A 1% inoculum was used for each serial transfer. Transfers were performed every 10 d. Error bars indicate SD, n=4.

**FIG S5.**
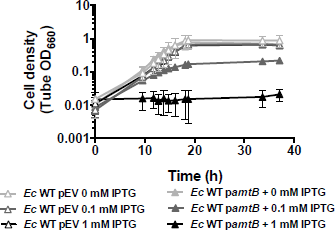
Increased *amtB* expression is harmful to *E. coli* in monocultures with 15 mM NH_4_^+^. Growth curves of WT *E. coli* monocultures harboring a plasmid encoding an IPTG-inducible copy of *amtB (pamtB*, filled) or empty vector (pEV, open) and grown at different IPTG concentrations. All monocultures were supplemented with 15 mM NH_4_C1 and 5 µg/ml chloramphenicol to maintain the plasmid. Cultures were inoculated with a 1% inoculum from stationary monocultures grown in 0 mM IPTG. After inoculation, IPTG was added to the indicated final concentration. Error bars indicate SD, n=3. ND, not determined.

### Increased NH_4_^+^ cross-feeding levels can compensate for the absence of a nitrogen starvation response

The NtrC regulon is critical during periods of nitrogen starvation, activating a wide variety of genes that are important for scavenging diverse nitrogen sources (20). We hypothesized that higher *R. palustris* NH_4_^+^ cross-feeding levels could mitigate the poor growth of *E. coli* ΔNtrC in coculture by making the nitrogen starvation response less important for survival. Previously, we engineered an *R. palustris* Nx strain that excretes 3-times more NH_4_^+^ by deleting *R. palustris* NH_4_^+^ transporters encoded by *amtB1* and *amtB2* (NxΔAmtB) (10). N_2_-fixing bacteria use AmtB to reacquire NH_4_^+^ that leaks outside the cell, and ΔAmtB mutants thus accumulate NH_4_^+^ into the supernatant (10, 12, 21). In agreement with our hypothesis, cocultures with *R. palustris* NxΔAmtB exhibited similar growth trends regardless of the *E. coli* strain used (Fig. 4B, D). As *R. palustris* NxΔAmtB excretes more NH_4_^+^ than *R. palustris* Nx, it was previously shown to result in faster WT *E. coli* growth and subsequent fermentation rates in coculture, ultimately leading to the accumulation of consumable organic acids (Fig. S4B) and acidification of the medium, inhibiting *R. palustris* growth (10). Cocultures pairing *R. palustris* NxΔAmtB and *E. coli* ΔNtrC similarly exhibited growth (Fig. 4B,D), and fermentation profile trends (Fig. S4B) that were indistinguishable from cocultures pairing *R. palustris* NxΔAmtB with WT *E. coli*. These similar trends indicate that high *R. palustris* NH_4_^+^ excretion can eliminate the trends observed when the *E. coli* nitrogen starvation response is compromised due to a ΔNtrC mutation.

One possibility for why high NH_4_^+^ cross-feeding levels eliminate the need for *E. coli ntrC* is that the free NH_4_^+^ levels might be sufficiently high enough to prevent activation of the *E. coli* NtrC regulon. However, comparative RNA-seq and proteomic analyses revealed that the same *E. coli* genes within the NtrC regulon that were highly upregulated in cocultures pairing WT *E. coli* with *R. palustris* Nx were also upregulated in cocultures with *R. palustris* NxΔAmtB (Table 1, Table 2). Thus, even though the *E. coli* nitrogen-starvation response is activated when cocultured with *R. palustris* NxΔAmtB, this response is likely dispensable if there is sufficiently high NH_4_^+^ cross-feeding.

### *E. coli* NtrC is required for adequate AmtB expression to access cross-fed NH_4_^+^ in coculture

While a high level of *R. palustris* NH_4_^+^ excretion can compensate for an improper *E. coli* nitrogen-starvation response, less NH_4_^+^ excretion could potentially exaggerate problems emerging from the absence of NtrC. We previously constructed an *R. palustris* ΔAmtB strain that excreted 1/3^^rd^^ of the NH_4_^+^ than *R. palustris* Nx in monoculture and which could not coexist in coculture with *E. coli* ΔAmtB (12). The reason for this lack of coexistence was due to *R. palustris* ΔAmtB outcompeting *E. coli* ΔAmtB for the lower level of transiently available NH**4**+, thus limiting *E. coli* growth and thereby the reciprocal supply of fermentation products to *R. palustris* (12). Expression of *E. coli amtB* is thus important in coculture in order to maintain coexistence. Indeed, RNA-seq and proteomic analyses revealed that *E. coli* AmtB transcript and protein levels were upregulated in all cocultures pairing WT *E. coli* with any of the three *R. palustris* strains (Nx, NxΔAmtB, ΔAmtB) (Table 1, Table 2). We thus wondered whether *E. coli* ΔNtrC would coexist with the low NH_4+_-excreting strain *R. palustris* ΔAmtB in coculture, as *E. coli amtB* expression is transcriptionally activated by NtrC. Consistent with our previous findings, *R. palustris* ΔAmtB supported a high relative WT *E. coli* population in coculture (Fig. 4D) (12). When cocultured with WT *E. coli, R. palustris* ΔAmtB responds to NH_4_^+^ loss to *E. coli* by upregulating nitrogenase activity since it has a wildtype copy of NifA (12). As a result, *R. palustris* ΔAmtB cross-feeds enough NH_4_^+^ to stimulate a high WT *E. coli* frequency and subsequent accumulation of consumable organic acids, similar to cocultures with *R. palustris* NxΔAmtB (Fig 3D, Fig. S4B) (12). In contrast, when we paired *E. coli* ΔNtrC with *R. palustris* ΔAmtB, little to no coculture growth was observed (Fig. 4C), similar to previous observations in cocultures pairing *E. coli* ΔAmtB with *R. palustris* ΔAmtB (12). Cocultures inoculated with single colonies of each species in this pairing grew to low cell densities Fig. S4A), and cocultures inoculated from these cocultures resulted in little to no growth, even after prolonged incubation (Fig. 4C).

As AmtB is under the control of NtrC (20), we hypothesized that cocultures pairing *E. coli* ΔNtrC with *R. palustris* ΔAmtB resulted in insufficient *E. coli amtB* expression, leading to a decreased ability to capture NH**4**+, which *R. palustris* will reaquire if given the chance (12). We thus predicted that increased expression of *amtB* in *E. coli* ΔNtrC would result in increased net growth of both species, as *E. coli* ΔNtrC would be more competitive for essential NH_4_^+^ and be able to grow and produce more organic acids for *R. palustris* ΔAmtB. To test this prediction, we obtained a plasmid harboring an IPTG-inducible copy of *amtB (pamtB)* for use in *E. coli* ΔNtrC. AmtB is typically tightly regulated and only expressed when NH_4_^+^ concentrations are below 20 µM, as cells acquire sufficient NH_4_^+^ through passive diffusion of NH_3_ across the membrane at higher concentrations (22). Additionally, excessive NH_4_^+^ uptake through AmtB transporters that exceeds the rate of assimilation can result in a futile cycle, as excess NH_3_ inevitably diffuses outside the cell (19). We first tested the effect of pamtB in WT *E. coli* monocultures with 15 mM NH**4**CL Induction with 1 mM IPTG prevented growth whereas 0.1 mM IPTG permitted growth albeit at a decreased growth rate (Fig. S5). We thus decided to use 0.1 mM IPTG to induce *amtB* expression in all cocultures described below. In cocultures pairing *E. coli* ΔNtrC *pamtB* with *R. palustris* ΔAmtB, more growth was observed than in cocultures with *E. coli* ΔNtrC harboring an empty vector (pEV) (Fig. 6A). In cocultures with *E. coli* ΔNtrC pEV, the *R. palustris* ΔAmtB cell density increased whereas the *E. coli* cell density did not (Fig. 6B). The *R. palustris* growth was likely due to growth-independent cross-feeding of fermentation products from *E. coli* maintenance metabolism, a phenomenon we described previously (11). In contrast, cell densities of both species increased in cocultures pairing *R. palustris* ΔAmtB with *E. coli* ΔNtrC *pamtB* (Fig. 6C), in agreement with our hypothesis that poor *E. coli amtB* expression contributed to the lack of growth in this coculture pairing. While *E. coli amtB* expression in this coculture pairing was sufficient to restore growth of both species, there are likely other genes within the NtrC regulon that contribute to *E. coli* growth in coculture. For example, the *E. coli* NtrC-regulated serine/threonine kinase yeaG has been shown to play a role in survival during nitrogen starvation by promoting metabolic heterogeneity (23). Indeed, *E. coli* yeaG and its associated protein of unknown function yeaH are both highly upregulated in coculture (Table 1). Thus, while we cannot rule out that other genes within the *E. coli ntrC* regulon are not important for coculture growth, the necessity of NtrC to upregulate *amtB* is clearly important.

**FIG 6.**
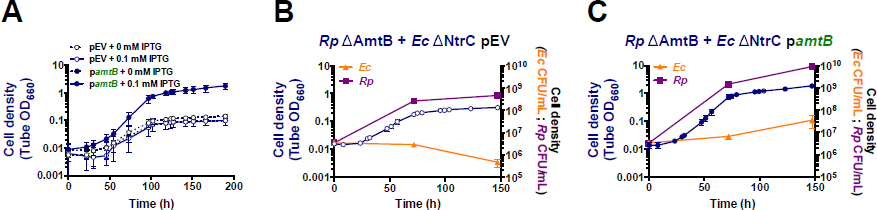
Ectopic expression of amtB in *E. coli* ΔNtrC permits mutualistic growth with *R. palustris* ΔAmtB. Growth curves (**A-C**) and cell densities for each species (**B, C**) from cocultures pairing *R. palustris (Rp)* ΔAmtB with *E. coli (Ec)* ΔNtrC harboring a plasmid encoding an IPTG-inducible copy of *amtB (pamtB*, filled circles) or an empty vector (pEV, open circles). To maintain plasmids, all cocultures were supplemented with 5 jrg/ml chloramphenicol, which is otherwise lethal to *E. coli* but not to *R. palustris* (Fig. S6). Cocultures were inoculated with a single colony of each species (**A**) or at a 1:1 starting species ratio based on equivalent CFUs/mL from starter *R. palustris* and *E. coli* monocultures (**B, C**). 0.1 mM IPTG was added to the cocultures at the initial time point. Error bars indicate SD, n=3.

**FIG S6.**
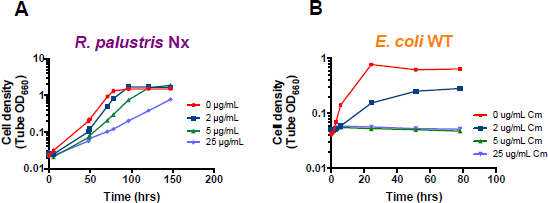
Determination of a chloramphenicol concentration to maintain pamtB in *E. coli* without harming *R. palustris*. Representative growth curves of *R. palustris* Nx (**A**) and WT *E. coli* (**B**) at different concentrations of chloramphenicol. All cultures were grown anaerobically in MDC with a 1% inoculum from stationary monocultures. *R. palustris* Nx was provided 20 mM sodium acetate as a carbon source with a 100% N_2_ headspace for nitrogen. WT *E. coli* was provided 25 mM glucose, 10 mM cation solution, and 15 mM NH_4_Cl.

## Discussion

In this study, we found that reciprocal nutrient cross-feeding between *E. coli* and *R. palustris* resulted in significant changes in gene expression in both species compared to monocultures. Based on the RNA-seq and proteomic analyses, we determined that *E. coli* alters its physiology to adopt a nitrogen-starved state in response to low NH_4_^+^ cross-feeding levels from *R. palustris*. We subsequently determined that this nitrogen-starved state is important for coexistence as genetic elimination of the master transcriptional regulator, NtrC, resulted in variable population outcomes. Mutualistic nutrient cross-feeding has also been shown to change the lifestyle of interacting partners in other systems. In natural communities, nutrient cross-feeding can alter gene-expression patterns to adapt each species to a syntrophic lifestyle (24–27). In some cases, the lifestyles exhibited within a mutualism might not even be possible during growth in isolation. For example, in synthetic communities that pair the sulfate-reducer *Desulfovibiro vulgaris* with the methanogen *Methanococcus maripaludis*, the methanogen consumes H_2_, which maintains low partial pressures that permit the sulfate reducer to adopt a fermentative lifestyle that would otherwise be thermodynamically infeasible (5). Similarly, in an experimental *Geobacter* coculture, direct electron transfer from *Geobacter metallireducens* to *Geobacter sulfurreducens* makes ethanol fermentation by *G. metallireducens* thermodynamically possible (7).

Similar to our mutualistic system, the mutualism between *D. vulgaris* and *M. maripaludis* represents a facultative mutualism, at least in the short term prior to evolutionary erosion of independent lifestyles (28). For mutualistic relationships to persist between partners that are conditionally capable of a free-living lifestyle, the relationship must exhibit resilience, or the ability to recover its function after a disturbance (29). One important resilience factor is the activation of regulatory networks that allow for microbes to quickly respond to environmental perturbations. Whereas flexible gene expression is useful for an individual microbe’s survival, excessive flexibility can sometimes lead to community collapse between mutualists in a fluctuating environment (30, 31). In the coculture of *D. vulgaris* and *M. maripaludis*, alternating between coculture and monoculture conditions, which require different metabolic lifestyles, resulted in community collapse (30, 31). Surprisingly, community collapse could be avoided by mutations that disrupted the *D. vulgaris* regulatory response needed to adapt cells for optimal growth rates in monoculture (30). Disruption of this regulatory response resulted in a heterogeneous *D. vulgaris* population, ensuring that a subpopulation would be primed for immediate mutualistic growth upon transition between growth conditions (31). In our system, the *E. coli* nitrogen starvation regulatory network was specifically activated by coculturing with *R. palustris* and was important for coculture stability. It is currently unclear if transitioning *E. coli* between monoculture and coculture conditions would result in similar community collapse or whether the NtrC-regulated network would adjust rapidly enough to meet the demands of each condition.

Nutrient starvation and other stress responses are widely conserved in diverse microbes and are primarily regarded as necessary for an individual’s survival in nutrient-limited environments (32–35). Many microbial communities are composed of primarily slow-growing or even non-growing subpopulations (36–38). However, lack of microbial growth in these communities does not imply cessation of cross-feeding, as bacteria often carry out growth-independent maintenance processes at slow rates (39), and such activities can be coupled to cross-feeding (11). Our findings suggest that nutrient starvation and perhaps other stress responses can help stabilize microbial cross-feeding interactions, especially at low nutrient cross-feeding levels. The extent to which specific starvation or stress responses are active in diverse mutualistic relationships remains unclear, yet likely depends on the environmental context. Together our results highlight the important role that alternate physiological states, including stress responses, can play in establishing and maintaining mutualistic cross-feeding relationships.

## Materials and Methods

### Strains and growth conditions

Strains, plasmids, and primers are listed in Table S1. All *R. palustris* strains contained *ΔuppE* and *ΔhupS* mutations to facilitate accurate colony forming unit (CFU) measurements by preventing cell aggregation (40) and to prevent H_2_ uptake, respectively. *E. coli* was cultivated on Luria-Burtani (LB) agar and *R. palustris* on defined mineral (PM) (41) agar with 10 mM succinate. (NH_4_)_2_SO_4_ was omitted from PM agar for determining *R. palustris* CFUs. Monocultures and cocultures were grown in 10 mL of defined M9-derived coculture medium (MDC) (10) in 27-mL anaerobic test tubes under 100% N_2_ as described (10). For harvesting RNA and protein, 100-mL cultures were grown in 260-mL serum vials. In both cases, MDC was supplemented with cation solution (1 % v/v; 100 mM MgSO_4_ and 10 mM CaCl_2_) and glucose (25 mM), unless indicated otherwise. *R. palustris* monocultures were further supplemented with 15 mM sodium bicarbonate, 7.8 mM sodium acetate, 8.7 mM disodium succinate, 1.5 mM sodium lactate, 0.3 mM sodium formate, and 6.7mM ethanol. *E. coli* monoculturas were further supplemented with 2.5 mM NH**4**CL Kanamycin was added to a final concentration of 30 µg/ml for *E. coli* where appropriate. Chloramphenicol was added to a final concentration of 5 µg/ml for both *R. palustris* and *E. coli* where appropriate. All cultures were grown at 30°C laying horizontally under a 60 W incandescent bulb with shaking at 150 rpm. Starter cocultures were inoculated with 200 µL MDC containing a suspension of a single colony of each species. Test cocultures and serial transfers were inoculated using a 1% dilution from starter cocultures. For experiments requiring a starting species ratio of 1:1, *E. coli* and *R. palustris* starter monocultures were grown to equivalent cell densities, and inoculated at equal volumes.

**Table S1.**
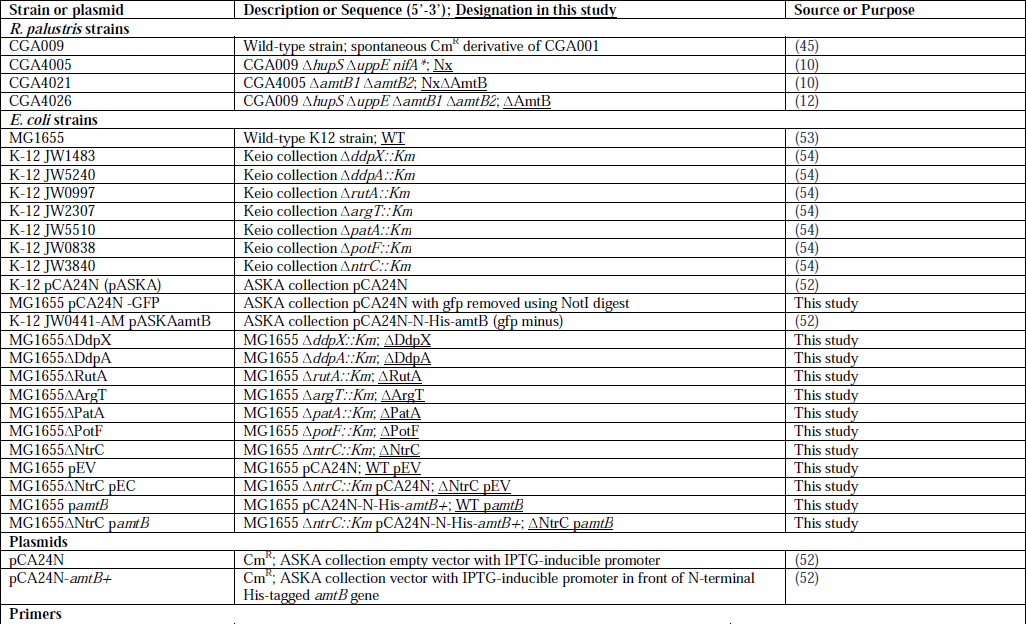
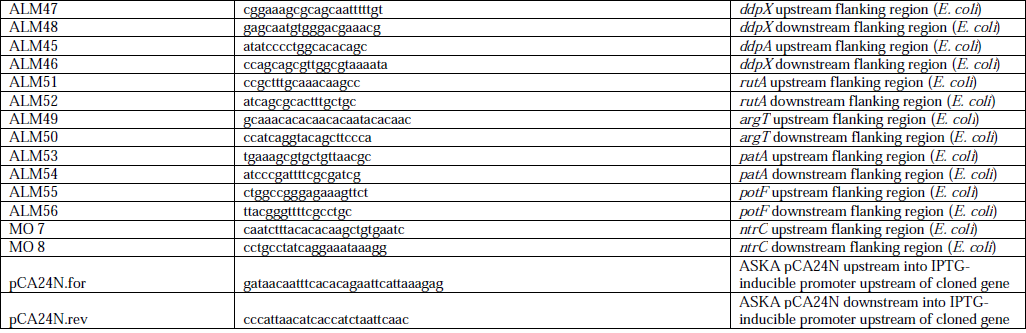
Strains and plasmids

### Generation of *E. coli* mutants

P1 transduction (42) was used to introduce deletions from Keio collection strains into MG1655. The genotype of kanamycin-resistant colonies was confirmed by PCR and sequencing.

### Analytical procedures

Cell density was assayed by optical density at 660 nm (OD_660_) using a Genesys 20 visible spectrophotometer (Thermo-Fisher, Waltham, MA, USA). Growth curve readings were taken in culture tubes without sampling (i.e., tube OD_660_). Specific growth rates were determined using readings between 0.1-1.0 OD_660_ where there is linear correlation between cell density and OD_660_. Final OD_660_ measurements were taken in cuvettes and samples were diluted into the linear range as necessary. Glucose, organic acids, formate and ethanol were quantified using a Shimadzu high-performance liquid chromatograph (HPLC) as described (43).

### Sample collection for transcriptomics and proteomics

Monocultures and cocultures were grown in 100-mL volumes to late exponential phase and chilled in an ice-water bath. A 1-mL sample was collected for protein quantification using a Pierce BCA Protein Assay Kit as per the manufacturer’s protocol. A 5- ml sample was removed for RNA extraction and 90 ml was used for proteomic analysis. All samples were centrifuged at 4°C, supernatants discarded, and cell pellets frozen in liquid N_2_ and stored at −80°C.

### RNA-seq

Total RNA was isolated from cell pellets using the RNeasy kit (Qiagen, Valencia, CA, USA) as per the manufacturer’s protocol. In order to calculate baseline expression levels, RNA sequencing reads resulting from monoculture were mapped to their corresponding reference genome *(E. coli* str. K-12 substr. MG1655 (44), NCBI RefSeq: NC_000913.3; *R. palustris* CGA0009 (45), NCBI RefSeq: NC_005296.1) using the Tuxedo protocol for RNA expression analysis (46) (Workflow deposited at https://github.com/behrimg/Task3/RNASeq). Specifically, split-reads were aligned to the reference genome with Tophat2 (v.2.1.0) (47) and Bowtie2 (v.2.1.0) (48). Following mapping, transcripts were assembled with cufflinks (v.2.2.0) (49), and differential expression was identified with the cufflinks tool, cuffdiff (v.2.2.0). To assure that crossmapping of homologous sequencing reads would not complicate expression analysis from the co-culture experiments, monoculture reads were additionally mapped as described to the opposing genome. As all potential crossmapping was confined to residual rRNA reads, these regions were excluded from the analysis and the co-culture RNA-seq reads where analyzed by mapping the sequenced reads to both reference genomes with no further correction.

### Preparation of protein samples for MS

Cell pellets were resuspended in 1 mL total protein buffer (TPB; 20mM HEPES-NaOH pH7.4, 150mM NaCl, 2mM EDTA, 0.2mM DTT, 1:100 PMSF, 1:100 protease inhibitors cocktail IV) and sonicated at 20% intensity (7 seconds on, 7 seconds off) for 5 min in an ice bath. Then 1/10 volume of 20% SDS was added. Samples were vortexed, boiled for 5 min, and immediately placed on ice. Debris was cleared by centrifuging for 30 s at 10,000 × g at 4°C and the supernatant was collected. Protein content of different lysates was analyzed by Coomassie staining following SDS-PAGE and sample aliquots containing 200 µg protein were subjected to chloroform:methanol protein extraction as described (50).

### Analysis by LC-MS/MS

Mass spectrometry was performed at the Mass Spectrometry and Proteomics Research Laboratory (MSPRL), FAS Division of Science, at Harvard University. Samples were individually labeled with tandem mass tag (TMT) 10-plex reagents according to the manufacturer’s protocol (ThermoFisher Scientific) and mixed. The mixed sample was dried in a speedvac and re-diluted with Buffer A (0.1 % formic acid in water) for injection for HPLC runs. The sample was submitted for a single liquid chromatography coupled to tandem mass spectrometry (LC-MS/MS) experiment which was performed on a LTQ Orbitrap Elite (ThermoFisher Scientific) equipped with Waters (Milford, MA) NanoAcquity HPLC pump Peptides were separated onto a 100 µm inner diameter microcapillary trapping column packed first with approximately 5 cm of C18 Reprosil resin (5 µm, 100 Å, Dr. Maisch GmbH, Germany) followed by analytical column ∼20 cm of Reprosil resin (1.8 µm, 200 Å, Dr. Maisch GmbH, Germany). Separation was achieved through applying a gradient from 5—27% ACN in 0.1% formic acid over 90 min at 200 nl min-1. Electrospray ionization was enabled through applying a voltage of 1.8 kV using a home-made electrode junction at the end of the microcapillary column and sprayed from fused silica pico tips (New Objective, MA). The LTQ Orbitrap Elite was operated in data-dependent mode for the mass spectrometry methods. The mass spectrometry survey scan was performed in the Orbitrap in the range of 395 —1,800 m/z at a resolution of 6 × 10^4^, followed by the selection of the twenty most intense ions (TOP20) for CID-MS2 fragmentation in the ion trap using a precursor isolation width window of 2 m/z, AGC setting of 10,000, and a maximum ion accumulation of 200 ms. Singly charged ion species were not subjected to CID fragmentation. Normalized collision energy was set to 35 V and an activation time of 10 ms. Ions in a 10 ppm m/z window around ions selected for MS2 were excluded from further selection for fragmentation for 60 s. The same TOP20 ions were subjected to HCD MS2 event in Orbitrap part of the instrument. The fragment ion isolation width was set to 0.7 m/z, AGC was set to 50,000, the maximum ion time was 200 ms, normalized collision energy was set to 27V and an activation time of 1 ms for each HCD MS2 scan.

### Mass spectrometry data analysis

Raw data were submitted for analysis in MaxQuant 1.5.**6**.5 (13). Assignment of MS/MS spectra was performed by searching the data against a protein sequence database including all entries from the *E. coli* MG1655 proteome (51), the *R. palustris* CGA009 proteome (45), and other known contaminants such as human keratins and common lab contaminants. MaxQuant searches were performed using a **20** ppm precursor ion tolerance with a requirement that each peptide had N termini consistent with trypsin protease cleavage, allowing up to two missed cleavage sites. 10-plex TMT tags on peptide amino termini and lysine residues were set as static modifications while methionine oxidation and deamidation of asparagine and glutamine residues were set as variable modifications. MS2 spectra were assigned with a false discovery rate (FDR) of 1% at the protein level by target-decoy database search. Per-peptide reporter ion intensities were exported from MaxQuant (evidence.txt). Only peptides with a parent ion fraction greater than or equal to 0.5 were used for subsequent analysis (6063 of 9987 peptides). Intensities were calculated as the sum of peptide intensities. Ratios between conditions were computed at the peptide level, and the protein ratio was computed as the mean of peptide ratios. All ratios were normalized by dividing by the median value for proteins from the same species. Ratio significance for coculture conditions at an FDR of 1% was computed by determining the ratio *r* at which 99% of genes have ratio less than *r* when comparing biological replicate monocultures.

### Expression of *E. coli amtB* in coculture

The ASKA collection (52) plasmid harboring an IPTG-inducible copy of *amtB* (pCA24N *amtB)* was purified from strain JW0441-AM and introduced by electroporation into WT *E. coli* and *E. coli* ΔNtrC. Cocultures were inoculated with either single colonies of each species or at a 1:1 starting species ratio, as indicated in the figure legends. IPTG and 5 µg/ml chloramphenicol were supplemented to cocultures to induce *E. coli amtB* expression in cocultures and maintain the plasmid, respectively.

## Acknowledgments

We thank B. A. Budnik and R. A. Robins (Harvard MSPRL) for assistance with mass spectrometry.

We thank P. L. Foster for providing the Keio and ASKA *E. coli* collections. This work was supported in part by the U.S. Department of Energy, Office of Science, Office of Biological and Environmental Research under Award Number DE-SC0008131 to JBM, by the U.S. Army Research Office, grant W911NF-14-1-0411 to ML, DAD, and JBM, by a National Institutes of Health National Service Award F32GM123703 to MGB, and by the Indiana University College of Arts and Sciences.

